# Nest Ecology-associated Impacts of Wastewater on Wild Bee Microbiomes

**DOI:** 10.1101/2025.09.09.675205

**Authors:** Lyna Ngor, Evan Palmer-Young, Helen Vo, Ali Alkhunder, Viraj Verma, Quinn McFrederick

## Abstract

Anthropogenic pollution affects environments differently depending on proximity to pollution source, exposure route, and species ecology. Thus, understanding organism’s ecological role and exposure route to contaminants is central to assessing pollution impact. Treated municipal wastewater releases contaminants into waterways and alters microbial communities. Plants absorb contaminants and expose animals through foraging and nest-building activities. Nesting ecology differences of ground vs wood cavity-nesting bees offers insight into niche-specific susceptibility to pollution. Because contaminants bind to soil strongly, ground-nesting bees near wastewater are likely most impacted, while wood cavity-nesting bees likely less impacted since plants’ ability to uptake contaminants are species dependent. We compared gut microbiomes of directly exposed soil-nesting *Halictus ligatus* and indirectly exposed wood-nesting *Ceratina* spp. upstream/downstream of wastewater. We collected bees, flowers, and soil, and analyzed their bacteria microbiomes (16S rRNA). Wastewater altered ground-nesting *H. ligatus* microbiome >18 times greater than wood cavity-nesting *Ceratina* adults. *Ceratina* larvae and pollen provisions showed significant but smaller shifts. Conversely, soil and flower microbiomes remained stable, indicating higher resilience. These results demonstrate that exposure routes drive contaminants susceptibility, with animal-associated microbes most vulnerable. Because bees are important pollinators and biodiversity contributors, these disruptions point to broader ecological risks in increasingly contaminated landscapes.

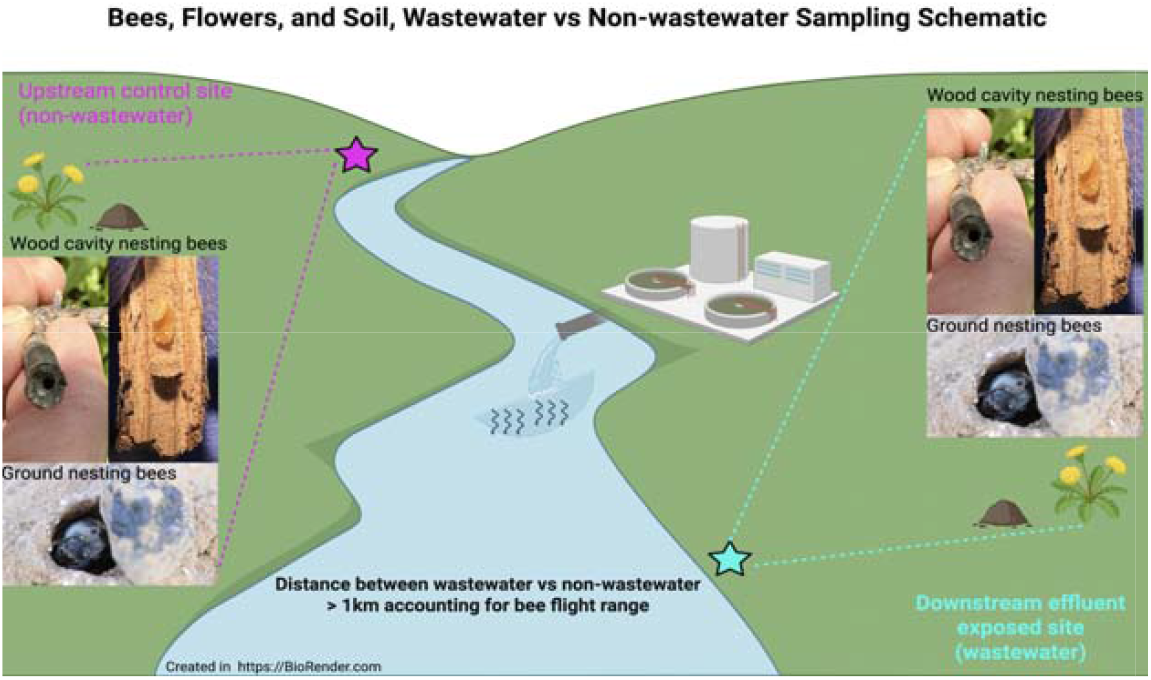

## Introduction

Urbanization, industrial activity, and agriculture all create pollution and pose a global threat to biodiversity and microbial ecology. Municipal wastewater, a major byproduct of urbanization, exemplifies how human activities introduce contaminants into natural systems. Although treatment plants remove many harmful substances, they often fail to eliminate effluent-associated contaminants, which persist in effluent and accumulate in the environment (1,2). These effluent contaminants can alter microbial diversity and function in both environmental and host-associated communities, disrupting ecological interactions and potentially compromising plant and animal health (3–6). Further, wastewater effluent creates nutrient rich environments that are optimal for certain bacteria taxa to exploit and proliferate (7–11) possibly leading to a competitive advantage to other taxa that may serve a role in supporting plant and insect health.

Beyond direct contamination of aquatic ecosystems, terrestrial reservoirs such as soil and flowering plants also harbor pollutants along with perturbed microbial communities and mediate exposure of animals to these compounds and their effects. Soil, particularly in urban areas receiving treated effluent, can accumulate contaminants (12) while simultaneously supporting complex microbial networks. Likewise, flowers serve as microbial hotspots, harboring diverse microbial communities that can be shaped by environmental conditions, including pollution (13). Floral-borne microbes form the microbiomes of many native bees (14) and are ecologically important via their effects on floral phenotypes and plant-pollinator interactions (15). However, little is known about how pollution-driven changes in soil and floral microbiota may cascade into higher trophic levels such as the microbial communities of native bees. Understanding how wastewater disrupts microbial communities is therefore critical to assessing the broader ecological implications of wastewater pollution.

Bees are ideal models to investigate how environmental pollution affects host microbiomes and how niche-specific susceptibility determines the severity of pollution exposure. Most native bees acquire their gut microbiota from environmental sources rather than through social transmission (16,17), making them especially vulnerable to pollution-induced shifts in soil and floral microbial communities such as in urban wastewater habitats. Microbiome dysbiosis can impair digestion, immunity, and overall fitness (16), while a stable microbiome has been shown to buffer bees against stressors like metal toxicity (18). Given native bees’ diverse nesting ecologies, their exposure to effluent contaminants vary: soil-nesting species, like *Halictus ligatus*, are in direct contact with contaminated soil and floral resources, while cavity-nesting species, like *Ceratina* spp., primarily experience indirect exposure through plant materials. These differences offer a unique opportunity to examine how pollution shapes microbiomes across environments and different host species’ ecologies.

In this study, we examine the effects of municipal wastewater pollution on microbial communities in native bees, soil, and flowers from sites up- or down-stream of wastewater treatment plants. By comparing a ground-nesting bee (*H. ligatus*) and a cavity-nesting bee (*Ceratina* spp.) at sites with and without wastewater exposure, we assess how bees’ niche-specific susceptibility and exposure routes shape bee-associated microbiota. We also characterize soil and floral microbial communities to understand how environmental reservoirs of microbes and contaminants correlate with bee microbiomes.

## Method

### Study-system

*Halictus ligatus* is a soil nesting bees that digs into the ground with their mouthparts to construct nests (19). Direct interaction with soil means that *H. ligatus* is likely exposed to effluent contaminants and its components in soil and groundwater in polluted areas. On the other hand, *Ceratina* spp. are wood cavity-nesting bees that excavate nests in pithy stems (20). Although *Ceratina* spp. do not come into direct contact with soil, they consume pollen, which may carry contaminants from soil that have been absorbed into plant tissues (21,22). By comparing the microbiomes of these two species in sites downstream versus upstream of wastewater treatment plants, we can determine if direct, indirect, or both direct and indirect exposure to effluent contaminants are potential drivers of bee-associated microbiota.

### Experimental Design

We collected bees, flowers, and soil at a total of 14 sites with two site types (see Map). 7 wastewater sites were downstream of wastewater effluent ejection into the natural river system, and non-wastewater sites comprise 7 locations that are upstream of any wastewater treatment facilities. Upstream reference sites were at least 1 km from wastewater outflows, based on bee flight range estimates relative to body size (23), to reduce overlap in foraging zones (supp. material: Table1).

### Map: Bees Field Collection Sites

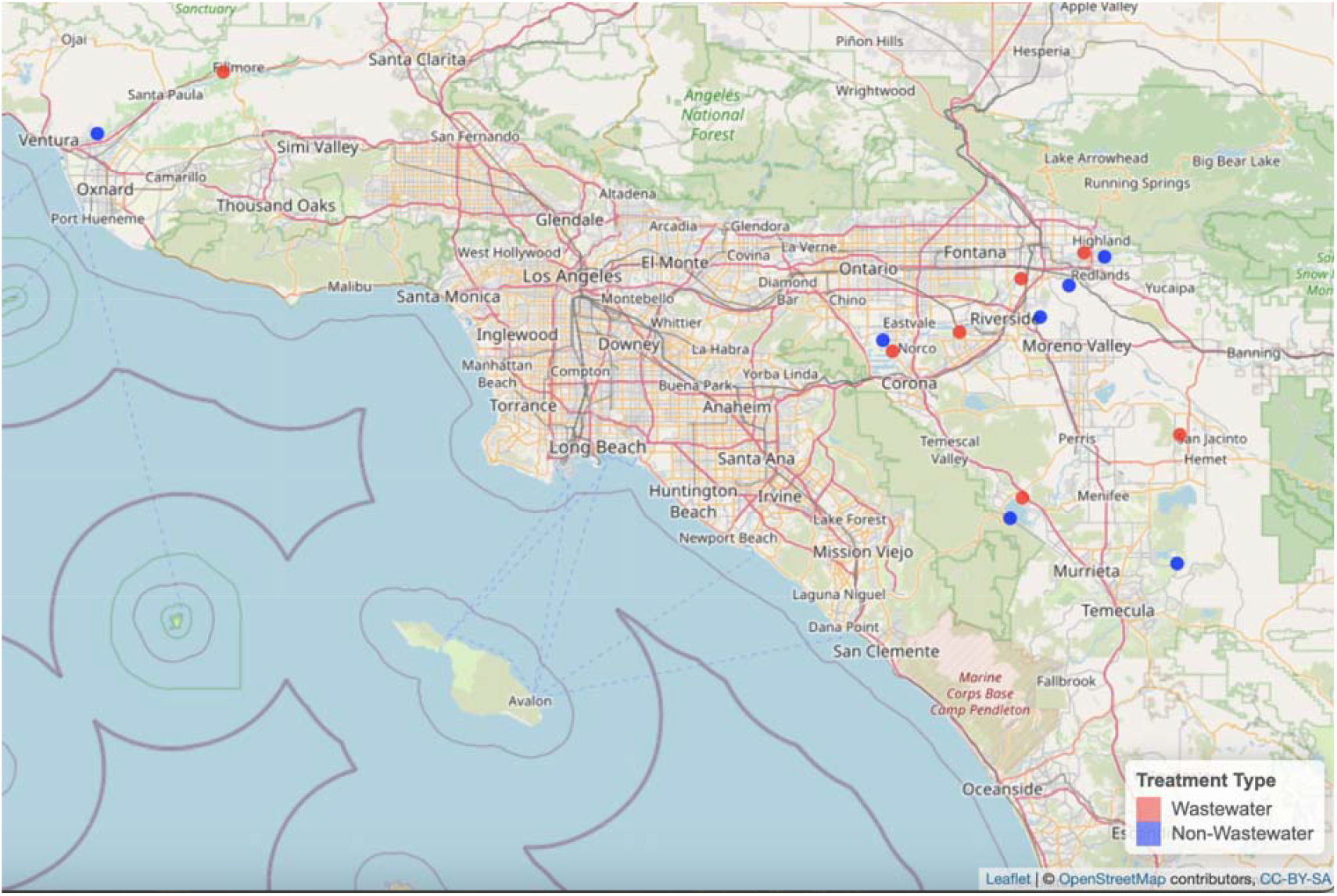

As underground nests are difficult to locate and excavate, we only collected foraging adults of *H. ligatus*. For *Ceratina* spp., we collected the whole nest with adults, larvae, and pollen provisions. Individual *H. ligatus* adults were stored in liquid nitrogen or dry ice during fieldwork and deposited in a −80C freezer until DNA extraction could be performed. *Ceratina* whole nests (in twigs) were stored in a cool ice box during field work. In the lab, *Ceratina* nests were aseptically dissected. Adults, larvae, and pollen provisions were separated into individual sterile tubes. Overall, we collected 4 types of host samples: *H. ligatus* adults, *Ceratina* spp. adults, *Ceratina* larvae, and *Ceratina* pollen provisions in all of our 14 sites. We analyzed *H. ligatus* and *Ceratina* spp. adult bees, *Ceratina* larvae and pollen provision independently. At each site, we collected ~ 50 mL of flowers on which we found *H. ligatus* foraging. We also collected elderberry tree (*Sambucus mexicana*) flowers from the same trees in which we found *Ceratina* nests. We haphazardly collected surface soil samples at each site into sterile 50 ml Falcon tubes.

#### Library preparation

We performed surface sterilization of *H. ligatus* adults, *Ceratina* adults, and *Ceratina* larvae by washing with 1% bleach (1 minute) followed by DI water twice (2 minutes). We performed whole gut dissection on *H. ligatus* adults, including the Dufour’s gland. Due to their small size, we used the whole abdomen of *Ceratina* adults and intact larvae. To obtain sufficient DNA for extraction, we pooled flower samples by host association and site. We added 10 ml of PBS per 50 ml Falcon tubes. We then agitated the tubes for 15 minutes using a Corning LSE vortex mixer at medium speed, followed by centrifugation at 18,000 rpm for 10 minutes to collect bacterial pellets. We used 1250 mg of soil material for each of the 14 sites. We conducted DNA extraction on adult bee guts and larvae using Qiagen’s Blood and Tissue kit, Qiagen Plant Pro kit for *Ceratina* pollen provision and flower wash, and Qiagen PowerSoil Pro kit for soil samples according to manufacturer’s protocol (Qiagen, Valencia, CA).

For bee guts, we lysed our samples using TissueLyser II Qiagen at 30 Hz for 6 minutes. Before lysing, we added 20ul ProK and 180 ul ATL buffer, two 3.2 mm steel beads, and ~100ul zirconia beads in each sample. After lysing, we incubated our samples for one hour at 56 °C. We added two negative controls that we carried through to all downstream steps. For flowers and soil, we follow the manufacturer’s protocol (Qiagen, Valencia, CA). Upon DNA quality verification, we prepared bees and flower wash samples for 16s Illumina barcoded by performing a two steps PCR as previously described (14). In PCR 1, we adapted each sample with its own unique primer sequences for downstream identification using the plant-plastid avoiding 799Fmod3 (CMGGATTAGATACCCKGG) and 1115R (AGGGTTGCGCTCGTTG) primers (see Table S1 for unique primer seq.). In PCR 2, we completed the Illumina sequencing construct. To ensure an equal molar proportion of each sample input across all samples, we performed PCR normalization using SequelPrep normalization plates (Invitrogen, Carlsbad, CA). Finally, samples were pooled and sent to University of California, Riverside Genomic Core for bioanalyzer 2100 (Aligent, Santa Clara, U.S.A.) and sequencing service using NextSeq 2000 P2 600 kit.

For soil samples, we processed extracted DNA for full length 16S rRNA gene sequencing using the Oxford Nanopore 16S Barcoding kit 24 V14 according to manufacturer’s protocol (Oxford Nanopore, Alameda, CA). As plant-plastid contamination is not problematic for soil samples, we were able to generate nearly full-length 16S rRNA gene sequences using 27F and 1492 primers. After the PCR protocol described by the manufacturer, we pooled the amplicons, loaded the MinION flow cell (FLO-MIN114), and sequenced using the MinION Mk1B sequencer. Before starting the sequencing device, we set the minimum qscore to 7 and the base calling model to fast basecalling.

#### qPCR H. ligatus adults

To determine if differences in microbiome structure were related to total bacterial load, we used qPCR to compare 16S rRNA gene copy number (absolute abundance) between wastewater and non-wastewater samples in the soil-dwelling *H. ligatus* adults. We performed qPCR reactions in triplicate using the same primers that we used for library preparation: 799Fmod3 and 1115R. Cycle times were converted to copy numbers using Gblock sequences (see supplemental materials) from 16s rRNA gene region for the bee-associated *Apilactobacillus* (24) as standard curves. A triplicate dilution series was included in the standard curve for all reactions. Amplification efficiency was between 90-110% in all of our reactions. See supplemental materials for qPCR protocol.

#### Bioinformatics Qiime2 for bees and flowers

We processed the 16S rRNA gene amplicons using the Qiime2 pipeline (25). We excluded any reads with a quality score below 35, and we binned exact amplicon sequence variants (ASVs) using DADA2 (26). We used the SILVIA 138 database (27) and QIIME2’s scikit-learn (28) to assign taxonomy to the Amplicon Sequence Variances (ASVs). We removed reads that were assigned to chloroplast and mitochondria from our dataset. We also eliminated any ASV that showed up only once (singletons). To avoid any potential reagent contaminant, we used decontam package in Qiime2 (25) to filter out any reads that showed up in our blank controls with a cutoff threshold of 0.5, which was 534 reads (0.001%) of the total reads (51,522,200 reads). In total, we retained 51,521,666 reads after quality controls for our downstream analysis.

#### Nanopore 16S amplicons using EPI2ME and Docker

We processed the 16S rRNA gene amplicon sequencing data using the epi2me-labs/wf-16s pipeline (v1.4.0) via Nextflow (v23.04.2) with Docker. Nanopore FASTQ files were quality filtered at a Q score threshold of 7, followed by reference database preparation using default SILVA and NCBI taxonomy databases. Taxonomic classification was performed with Minimap2, and microbial abundance tables and summary reports were generated. Negative controls yielded fewer than ten reads and were excluded from downstream analysis.

#### Alpha refraction, alpha and beta diversity, and Aldex2 Taxonomic Differential Abundance

In the bee-associated samples, we separated our data into 4 groups by tissue source: adult *H. ligatus*, adult *Ceratina* spp., *Ceratina* larvae, and *Ceratina* pollen provision. Using 7 qiime2, we rarefied each of these datasets and chose a sampling depth at a point in which the observed features in the rarefaction graph leveled off. Sampling depth was set for the following: *H. ligatus* = 9,000 reads, *Ceratina* adult = 9,000 reads, *Ceratina* larvae = 20,000 reads, and *Ceratina* pollen provision = 50,000 reads (Table1). We set the sampling depth for our flower and soil samples at 5,000 and 200,000 reads respectively.

We assessed alpha diversity (observed ASVs, Pielou’s evenness) using Kruskal–Wallis tests. For beta diversity, we computed Bray–Curtis dissimilarities from rarefied ASV tables and visualized patterns with NMDS using metaMDS() in the vegan R package (Oksanen et al., 2017). We used PERMANOVA via adonis (vegan) to test for significant differences in community composition between wastewater and non-wastewater sites for each sample type. To assess within-group variability, we applied beta dispersion tests using betadisper() and permutest() (vegan), which evaluate heterogeneity in Bray– Curtis distances within site types. We used Moran’s I to test spatial autocorrelation in Shannon diversity. Differential abundance was assessed using ALDEx2 (QIIME 2 plugin) with FDR < 0.1 (Jiang et al., 2019; Fernandes et al., 2013), which applies compositional modeling and Monte Carlo sampling to detect ASVs enriched by treatment.

## Results

### Quality Control

A total of 77 out of 584 samples (13.18%) were excluded due to low sequencing depth based on alpha rarefaction thresholds. Across all filtering steps, we removed 5.99 × 10^7^ reads, the vast majority of which were lost during rarefaction subsampling. An additional 1.58 × 10^5^ reads were removed due to contamination or classification as non-target sequences (e.g., blanks, chloroplasts, mitochondria). After filtering, we retained 1.27 × 10^7^ high-quality reads for analysis. ASV reads were separated by host types such as *H. ligatus, Ceratina* adults, larvae, and pollen provision (<u>S1 Table 1</u>).

### qPCR for H. ligatus

To assess whether wastewater exposure altered absolute bacterial load in *H. ligatus* adults, we quantified 16S rRNA gene copy number via qPCR (n= 118). A one-way ANOVA showed no significant difference between site types (Mean sq= 1.42 × 10^12^; F_1,393_= 2.87; p = 0.0911). A two-way ANOVA including site as a factor confirmed that site types were not significant (Mean sq=1.42 × 10^12^; F_1,380_ = 3.14; p = 0.0771), while site itself was (Mean sq=1.74 × 10^12^; F_13,380_ = 3.86, p < 0.001), indicating high site-level variation. The interaction model had the lowest AIC (–15,506.31) but explained little variance, suggesting overfitting. Overall, wastewater exposure did not significantly affect total bacterial load in *H. ligatus* (<u>S1 Table 2</u>).

### NMDS Ordination Analysis and Beta Diversity Analyses

Our beta diversity analyses revealed that the abdominal microbial communities of *H. ligatus* adults exhibited the strongest response associated with proximity to wastewater, with PERMANOVA explaining 24.74% of the variation in Bray–Curtis dissimilarities between site types (*R*^*2*^ = 0.2474, *p* < 0.001; Figure 1A; supp. material: Table 3). *Ceratina* spp. adults also showed significant but smaller treatment differences (*R*^*2*^ = 0.0135; *p* < 0.05; Figure 1A, S1 Table 3). Similarly, *Ceratina* larvae and pollen provision microbiomes differed by treatment, with *R*^*2*^ values of 0.0415 and 0.0403, respectively (Figure 1B–C, S1 Table 3). In contrast, no significant treatment differences were detected in flower and soil microbiomes (S1Table 3), *p* > 0.1, suggesting higher environmental resilience.

**Figure 1.**
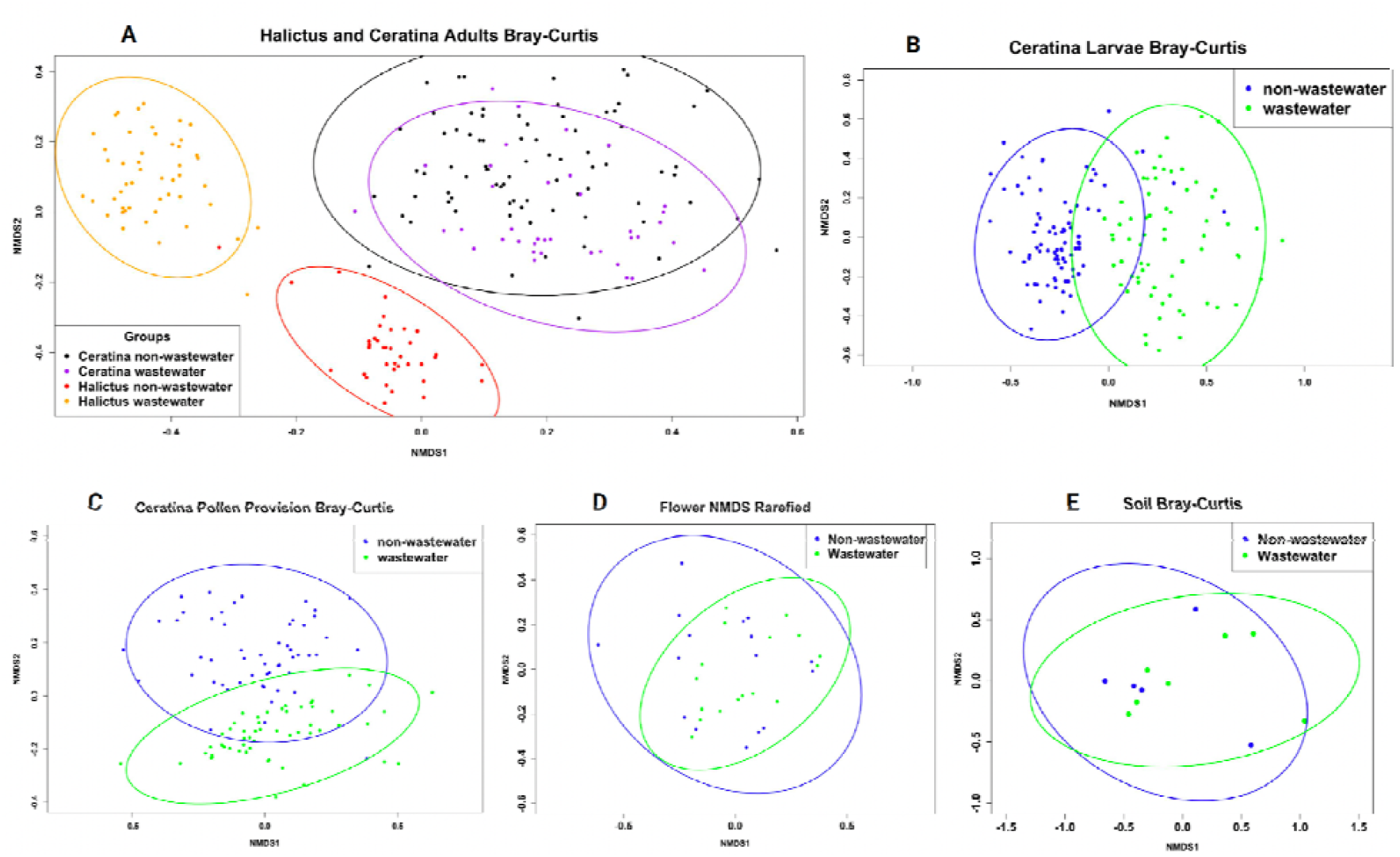
NMDS ordination of Bray–Curtis dissimilarities across wastewater and non-wastewater sites for microbial communities from bee, larvae, and pollen provisions showed distinct clusters while soils and flowers showed no distinction between site types. Each point represents an individual sample; ellipses indicate 95% confidence intervals for group centroids. **(A)** Gut microbiomes of adult *Halictus ligatus* and *Ceratina spp.* show strong separation by treatment in *H. ligatus (PERMANOVA* R^2^ = 0.2474, *p* < 0.001), and weaker but significant separation in *Ceratina* adults (R^2^ = 0.0135, *p* < 0.05). **(B)** *Ceratina* larval gut communities differ significantly between site types (R^2^ = 0.0415, *p* < 0.001), with moderate centroid separation. **(C)** Pollen provisions show modest but significant compositional differences (R^2^ = 0.0403, *p* < 0.001), suggesting environmental input from contaminated floral sources. **(D)** Floral surface microbial communities overlap extensively by treatment, with no significant differences (R^2^ = 0.026, *p* = 0.753). **(E)** Soil communities show no significant compositional shifts (R^2^ = 0.1208, *p* = 0.159), indicating relative stability under treated effluent exposure.

To complement these findings, we examined within-group variability using the betadisper() and permutest() functions in the vegan R package. Wastewater exposure significantly increased beta dispersion in *H. ligatus* adults (F_1,84_ = 34.85, *p* = 0.001;supp. material: Figure C1) and Ceratina pollen provisions (F_1,101_ = 10.09, *p* = 0.003; supp. material: Figure C4), indicating greater microbial heterogeneity under contamination. Ceratina larvae showed a marginal increase in dispersion (F_1,147_ = 3.99, *p* = 0.055; supp. material: Figure C3), while Ceratina adults (F_1,116_ = 2.82, *p* = 0.109; supp. material: Figure C2), floral washes (*p* = 0.91; supp. material: Figure C5), and soil samples (F_1,10_ = 2.11, *p* = 0.159; supp. material: Figure C6) did not differ significantly.

Together, these complementary analyses demonstrate that wastewater exposure not only shifts the average microbial composition (PERMANOVA) in sensitive hosts like *H. ligatus* and *Ceratina* brood stages, but also increases variability in microbial structure (beta dispersion), particularly in *H. ligatus* that are in close contact with contaminated soils. These findings emphasize both the directional and stochastic impacts of contamination on host-associated microbial communities.

### Alpha-diversity: Richness and Evenness

We assessed microbial alpha diversity using both Pielou’s evenness and species richness (observed ASVs) across adult H. ligatus, adult Ceratina spp., Ceratina larvae, pollen provisions, floral washes, and soil samples. All diversity metrics were calculated from rarefied ASV tables using QIIME 2 and analyzed with Wilcoxon rank-sum tests in R using the vegan, ggplot2, and ggpubr packages. Evenness differed significantly only in H. ligatus adults, with wastewater samples showing reduced community evenness (W = 1330, p < 0.001; Cliff’s delta = −0.45), suggesting dominance by a few taxa in wastewater treatment (supp. material: Figure D1). No significant differences in evenness were detected in *Ceratina* adults (W = 1309, p = 0.2254), larvae (W = 2403, p = 0.1899), pollen provisions (W = 1572, p = 0.5958), flower washes (p = 0.79), or soil (W = 25, p = 0.7789) (supp. material: Figure D2-D6). These results highlight a host-specific response in microbial balance under wastewater exposure, with *H. ligatus* being the most sensitive. Species richness remained stable across all sample types. No significant differences in observed ASVs were found between wastewater and non-wastewater sites for any host or environmental matrix (all p > 0.05; supp. material: Figure E1–E6). This indicates that while the relative abundance distribution may shift in *H. ligatus*, the overall number of microbial taxa remains unchanged.

### Aldex2 Differential Abundance

Our differential abundance analysis showed that *Enterococcus* was the most differentially abundant microbe in *H. ligatus* adults from wastewater environments, while *Wolbachia, Lactobacillus*, and *Gilliamella* dominated in non-wastewater environments (Figure 2A). In contrast, *Ceratina* adults showed *Wolbachia, Lactobacillus*, and *Gilliamella* were more abundant in wastewater environments (Figure 2B). *Ceratina* pollen provision samples had higher differential abundance of *Lactobacillus* and *Gilliamella* in non-wastewater habitats (Figure 2C). In *Ceratina* larvae, *Lactobacillus, Gilliamella*, and *Tyzzerella* were more abundant in non-wastewater environments (Figure 2D).

**Figure 2.**
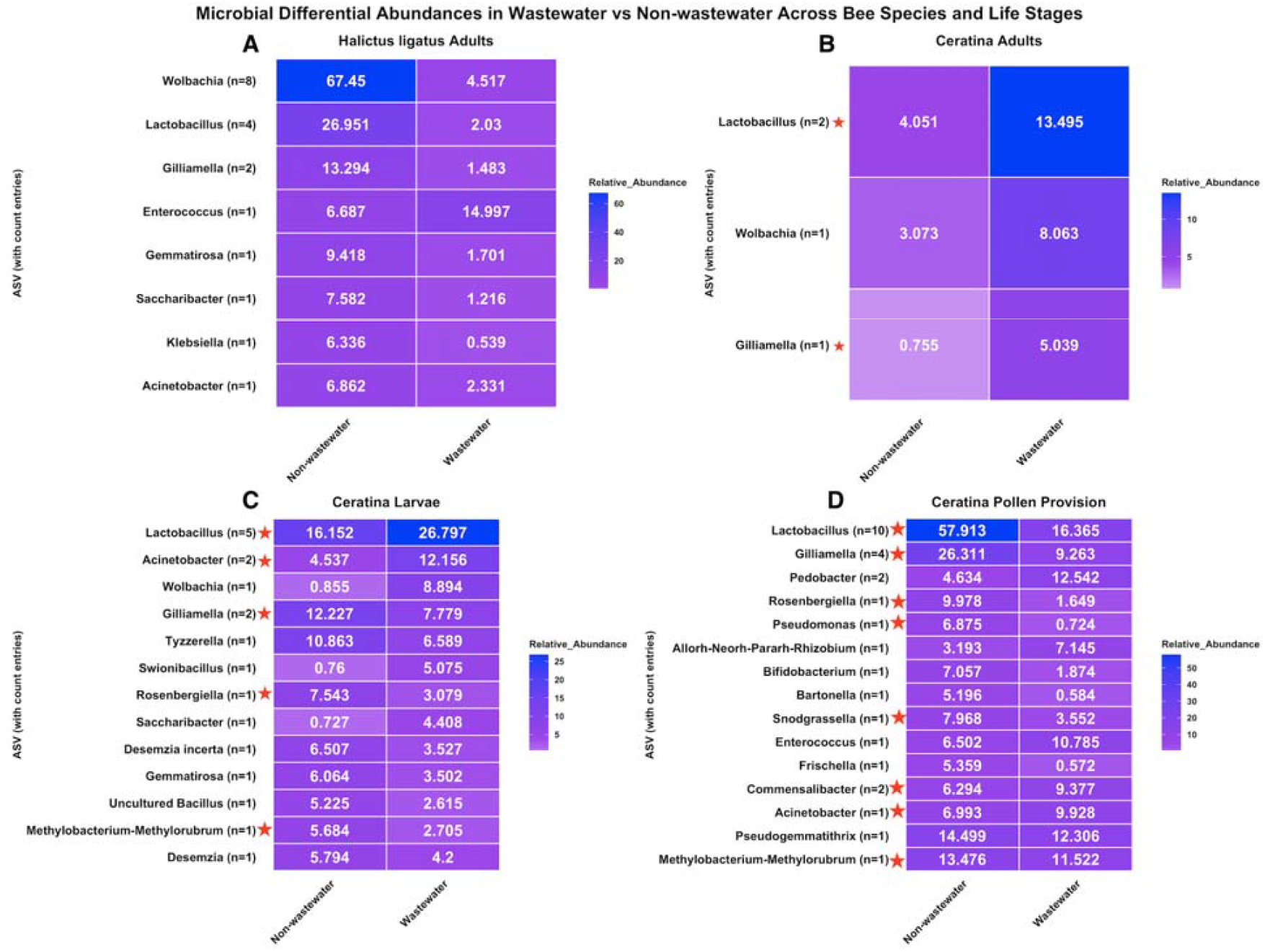
Heatmaps showing differentially abundant microbial genera in fold changes across the 4 sample types that showed a significant overall effect of proximity to wastewater: **(A)** *Halictus ligatus* adults, **(B)** *Ceratina* adults, **(C)** *Ceratina* larvae, and **(D)** *Ceratina* pollen provisions. Columns represent wastewater and non-wastewater samples. Each heatmap includes only genera with statistically significant differential abundance between site types. Color intensity reflects relative abundance, with deep blue indicating higher abundance and white indicating lower abundance. Numeric labels within each tile represent the summed relative abundance of each genus across all samples for that treatment group. Red star next to bacteria genera indicates *Ceratina* core taxa.

### Spatial Autocorrelation Moran I Using Beta Diversity Ordinations

We only conducted spatial autocorrelation for sample types that showed significant p-values <0.05 from our PERMANOVA result. Our analysis of spatial autocorrelation was assessed using Moran’s I for sample types with significant PERMANOVA results (p < 0.05; Supplemental Table 3). *Ceratina* larvae exhibited strong positive spatial autocorrelation (I = 0.4523, p < 0.001). In contrast, *Halictus ligatus* adults (I = –0.0613, p = 0.001) and *Ceratina* pollen provisions (I = –0.0339, p = 0.001) showed significant negative spatial autocorrelation. *Ceratina* adults did not display significant spatial structure (I = –0.0114, p = 0.175; Supplemental Figure B1).

## Discussion

Our comparison of bee-associated microbes between site types, with and without exposure to wastewater effluent indicates that microbes of ground-nesting bees (*Halictus ligatus*) showed the strongest association with wastewater exposure more than wood-cavity-nesting bees (*Ceratina* spp.). This result was driven by a major reduction in wild bee-associated microbes such as *Wolbachia, Lactobacillus*, and *Gilliamella* in *H. ligatus* and in *Ceratina* larvae and pollen provision that are exposed to wastewater effluent, while the same microbial taxa were elevated *Ceratina* adults exposed to wastewater effluent.

Further, we found that *Enterococcus* were highly abundant in wastewater-associated *H. ligatus* adult samples. This result likely reflected the ability of this bacteria to exploit the nutrient-rich, contaminated conditions typically found in polluted environments (7–11) and niche-specific susceptibility differences in each host. Effluent associated bacteria exhibit a variety of traits that promote survival in challenging conditions, such as resistance to high levels of organic pollutants, heavy metals, and antibiotics. Specifically, *Enterococcus* thrives in fecal-contaminated waters due to its ability to tolerate stress and acquire antibacterial genes from other microbes (29), while *Acinetobacter* is well known for its survival in a range of environments, including flowers, nectar, and hospital wastewater (30–32). In comparison, *Acinetobacter* has a wide range of antibiotic resistance capabilities (33) demonstrating their adaptability across diverse environments. These findings suggest that the microbial communities of municipal wastewater can have cascading effects on the microbiomes of wild animal hosts, with particular implications for the ground-nesting taxa that account for 70% of total native bee species in the U.S. (34).

The stark association between wastewater effluent exposure and *H. ligatus* gut microbiome structure could indicate that effluent contaminants are disrupting key symbiotic relationships critical to host health. Native bees acquire their gut microbiomes from the environment (35) including flowers, soil, and nest materials. Aside from exposure to pollution, flowers and soil are fundamentally different from each other, including in chemicals, pH, and temperature which play a major role in providing the condition for certain microbial taxa to thrive (36,37). Further, the differential resilience of symbiotic microbes to environmental pollutants (18) underscores broader ecological risks from pollution: the loss of core mutualists could diminish the functional capacity of bees to maintain health (38) and potentially reproductive success. The dominance of opportunistic and pollution-adapted taxa in wastewater habitats could additionally drive antibiotic resistance and alter nutrient cycling in pollinator-dependent ecosystems.

Although *Ceratina* adults, larvae, and pollen provision microbial communities showed statistically significant differences across site types (Figure 1; Supp. Table 1), their core taxa *Lactobacillus* and *Gilliamella* (39), remained dominant, and in adults, were even more abundant in wastewater-exposed individuals. This persistence of the core gut microbiome suggests that *Ceratina* microbial communities are more resilient to environmental disturbance, likely due to reduced, indirect exposure to soil-borne contaminants.

The contrasting association between wastewater exposure and *Wolbachia* abundance in *H. ligatus* versus Ceratina exemplifies the potential influence of nest ecology. *Wolbachia*, an endosymbiotic bacterium, can hijack host physiological function, and drive its own proliferation by manipulating sex ratios to favor infected females (40). While various strains and abundances of *Wolbachia* were found in many solitary bee species (39,41,42), they were not part of the *Ceratina* core gut microbiome. Notably, *Wolbachia* remained stable in *Ceratina* across wastewater vs non-wastewater sites, but were highly responsive in *H. ligatus*. Given Wolbachia’s known sensitivity to antibiotics (43), its decline in *H. ligatus* suggests that chemically enriched soils are driving shifts in both gut and intra-cellular symbiont composition that include this widespread and potentially phenotypically critical insect symbiont (40). These findings highlight divergent microbial responses based on host nesting ecology. The extent of exposure to contaminated materials appears to be the key risk factor: *Ceratina* nests in wood cavities, while *H. ligatus* directly contacts contaminated soil.

The movement of effluent-derived contaminants through the ecosystem results in a concentration gradient between waterways and other discharge sites, surrounding soils 14 and groundwater, and plant materials. Even though antibiotics and effluent contaminants can translocate into plant roots, stems, and leaves (44–46), their accumulation in floral tissues is minimal or undetectable (47). This limited floral contamination likely reduces effluent contaminants exposure in *Ceratina*, whose nesting and foraging behaviors minimize direct contact with contaminated substrates. In contrast, *H. ligatus* nests in sandy-clay mixes, firm silt, and hardened clay (19) that are rich in Al^3+^, Ca^2+^, and Fe^3+^. These soils have high cation exchange capacity, allowing strong binding of antibiotics (48). Additionally, in wastewater-impacted habitats, continuous input of effluent contaminants further enriches soils with substances that have strong antibiotic binding capacity, leading to chronic exposure for ground-nesting bees. This chemical exposure may favor antibiotic-resistant taxa like *Enterococcus,* which was dominant in *H. ligatus* guts but rare in soil, further supporting a chemically driven selection process.

Ceratina larvae and pollen provisions also showed statistically significant differences in relative abundance between wastewater and non-wastewater sites. *Ceratina* larvae core gut microbiome showed mostly similar patterns as in *Ceratina* adults affected by wastewater with a significant increase in abundance of *Lactobacillus* and *Acenitobacter*. On the contrary, *Gilliamella* was significantly reduced in wastewater-exposed *Ceratina* larvae (Figure 2C) while significantly increased in wastewater-exposed *Ceratina* adults. In comparison, *Ceratina* pollen provision exposed to wastewater showed a significant reduction in 8 taxa of their 13 core microbial taxa including *Lactobacillus, Gilliamella, Rosenbergiella, Pseudomonas, Snodgrassella, Commensalibacter, Acinetobacter, Mythylobacterium* (39) (Figure 2D), suggesting host physiology during larval development may modulate microbial colonization (24). These results suggest that while *Ceratina* adults may avoid direct environmental impact, the microbial communities associated with larvae and pollen provisions are more susceptible to wastewater exposure, with some core taxa demonstrating resilience while others show significant decline.

The differences in *Ceratina* adults, larvae, and pollen provision R^2^ values (supp. material: Table 2) from our PERMANOVA showed different susceptibilities across life stages of *Ceratina* bees. *Ceratina* pollen provisions and larvae were more affected by wastewater pollution than adults. This heightened vulnerability could be due to larvae’s direct exposure to contaminated pollen, which serves as their primary food source during development as shown by the declining abundance of one of its core taxa *Rosenvergiella. Ceratina* adults, on the other hand, demonstrated greater resilience, possibly due to more developed detoxification systems, a broader foraging range, and/or the stability of key microbial taxa such as *Lactobacillus* across wastewater and non-wastewater habitats (Figure 2). These differences emphasize the need to consider the entire life cycle when evaluating the ecological risks posed by contaminants, as microbes can affect development in some (49) but not all bees (50).

Although we observed no significant differences in microbial communities in flower surface washes and soil from wastewater vs non-wastewater sites, it is important to note that similar taxonomic profiles do not necessarily imply functional redundancy. Soil and flowers from wastewater and non-wastewater environments may host microbial communities with overlapping taxonomic identities yet diverge in their gene expression (51). Thus, functional shifts may still occur in the absence of clear taxonomic differentiation. Also, our surface layer soil sampling effort may have limited our ability to assess effluent contaminants impact on soil microbes. Future studies incorporating soil from greater depth and shotgun metagenomic sequencing may provide higher taxonomic resolution and more importantly, uncover functional genomic shifts. This approach could offer better insight into how wastewater influences not only microbial community composition but also potential functional consequences for native bees and surrounding ecosystems.

While the field of ecotoxicology has consistently demonstrated that wastewater effluent is impacting the microbial communities of nearby fauna, findings are often focused on organisms that lack distinctive niche-specific susceptibility between their own species (6,52). For instance, aquatic associated insects like mosquitoes are an insightful system in assessing wastewater effluent impact (3) with implication in disease vectors, but their ecological niche which determines their exposure to effluent contamination is similar between species. In contrast, our research leverages the distinct nesting habitats (soil-nesting versus wood cavity-nesting) of two related bee taxa, to associate the intensity of exposure to wastewater-contaminated substrates with the extent of wastewater-associated disturbance to the host microbiome. This pioneers the exploration of ecological niche-specific vulnerability to wastewater effluent exposure and its implications for native bees.

## Conclusion

Research in municipal wastewater effluent has primarily focused on the effects of effluent contaminants on specific study systems including soil, water, plants, and aquatic-associated animals ranging from mosquitoes to fishes. Here, we offer a new perspective. By leveraging niche-specific susceptibility in native bees, we ask how municipal wastewater effluent impacts the environment based on animal ecological niche. We present the first investigation into how wastewater effluent affects microbial communities differently in related host species but with distinct exposure to wastewater effluent due to their nesting strategies. Our results show that ground-nesting bees, particularly *Halictus* ligatus, are severely impacted, while cavity nesting *Ceratina* exhibit significant effects in larvae and pollen provision with a significant decline of one of its core taxa as well as variable responses across developmental stages. The bacterial taxa driving these changes differ between habitats, with wastewater-exposed environments dominated by wastewater-associated bacteria, and non-wastewater sites characterized by bee- and flower-associated bacteria.These findings highlight the potential for species-specific host ecology to moderate exposure to and microbial effects of a ubiquitous anthropogenic disturbance. Consideration of these differences could help to prioritize at-risk taxa in ecological risk assessments and conservation efforts.

## Acknowledgements

This work was supported by the National Science Foundation (NSF) award number 1929572 and USDA Hatch Funds CA-R-ENT-5109-H, both granted to Q.S. McFrederick. Additional support was provided by the Shipley Skinner Foundation, the Utom River Native American Conservation Funds Fellowship, and a California Institute of Biodiversity Grant.

We thank the University of California, Riverside Genomics Core for sequencing support and acknowledge the field and laboratory assistance provided by members of the McFrederick Lab. We thank Dr. Scott McArt, Dr. Wayne Anderson, Christina Zhao at Cornell Chemical Ecology Core Facility for soil chemistry analysis support. We are especially grateful to the park rangers and personnel who facilitated site access and sampling: Laura Pavliscak (Ventura Land Trust), Kalee Koeslag (Lake Skinner), and Dr. Jessica Maccaro, whose help with field access and sample collection was instrumental. We also thank Dr. Erin Rankin, Dr. Kurt Anderson, and Dr. Magda Guzman for their valuable guidance and input throughout the development of this project.

